# Protein products of non-stop mRNA disrupt nucleolar homeostasis

**DOI:** 10.1101/851741

**Authors:** Zoe H. Davis, Laura Mediani, Jonathan Vinet, Simon Alberti, Alex S. Holehouse, Serena Carra, Onn Brandman

## Abstract

Mutations that cause ribosome stalling or impair the cell’s protective response to stalling have been demonstrated to cause neurodegeneration, yet the mechanisms underlying these pathologies remain poorly understood. Here we investigated the fate of defective proteins translated from stall-inducing, nonstop mRNA that escape ubiquitylation by the Ribosome-associated Quality Control (RQC) E3 ligase LTN1. We found that nonstop protein products accumulated in nucleoli and this localization was driven by polylysine tracts produced by translation of the poly(A) tail of nonstop mRNA. Nucleolar sequestration increased the solubility of invading proteins but disrupted nucleoli, altering their dynamics, morphology, and resistance to stress. Changes in nucleolar morphology are consistent with a simple physical model in which LTN1 impairment enhances the inter-molecular interactions of nucleolar components. Our work elucidates how failure to degrade the protein products of stalled translation may affect distal cellular processes and will inform studies on the pathology of neurodegenerative disease.

## Introduction

Defects in mRNA can cause ribosomes to stall and abort translation(Doma and Parker, 2006; Frischmeyer et al., 2002a; Joazeiro, 2017). The most common mRNA defect is premature polyadenylation, in which the polyadenylation machinery utilizes cryptic signals upstream of the stop codon, placing a poly(A) tail within the open reading frame and truncating the mRNA (Frischmeyer et al., 2002a; van Hoof et al., 2002; Klauer and van Hoof, 2012; Tian et al., 2007). Premature adenylation upstream of the stop codon occurs in ~1% of all mRNAs (Frischmeyer et al., 2002a; Graber et al., 1999; Klauer and van Hoof, 2012; Zhou et al., 2018). Translation of these “nonstop” mRNAs proceeds into the poly(A) tail, where ribosomes translate several lysine-encoding AAA codons before stalling (Bengtson and Joazeiro, 2010a; Brandman et al., 2012a; Defenouillère et al., 2013a; Frischmeyer et al., 2002b; van Hoof et al., 2002; Ito-Harashima et al., 2007a; Juszkiewicz and Hegde, 2017; Sundaramoorthy et al., 2017; Verma et al., 2013). Premature polyadenylation therefore produces truncated nascent polypeptide chains appended with polylysine. To protect cells from these defective proteins, a process called Ribosome-associated Quality Control (RQC) detects stalled translation events and ubiquitylates stalled nascent chains via the E3 ubiquitin ligase Listerin (LTN1), targeting them for destruction by the proteasome (Bengtson and Joazeiro, 2010b; Brandman et al., 2012b; Defenouillère et al., 2013b; Ito-Harashima et al., 2007b; Shao et al., 2013) (Fig. 1a).

**Figure 1.**
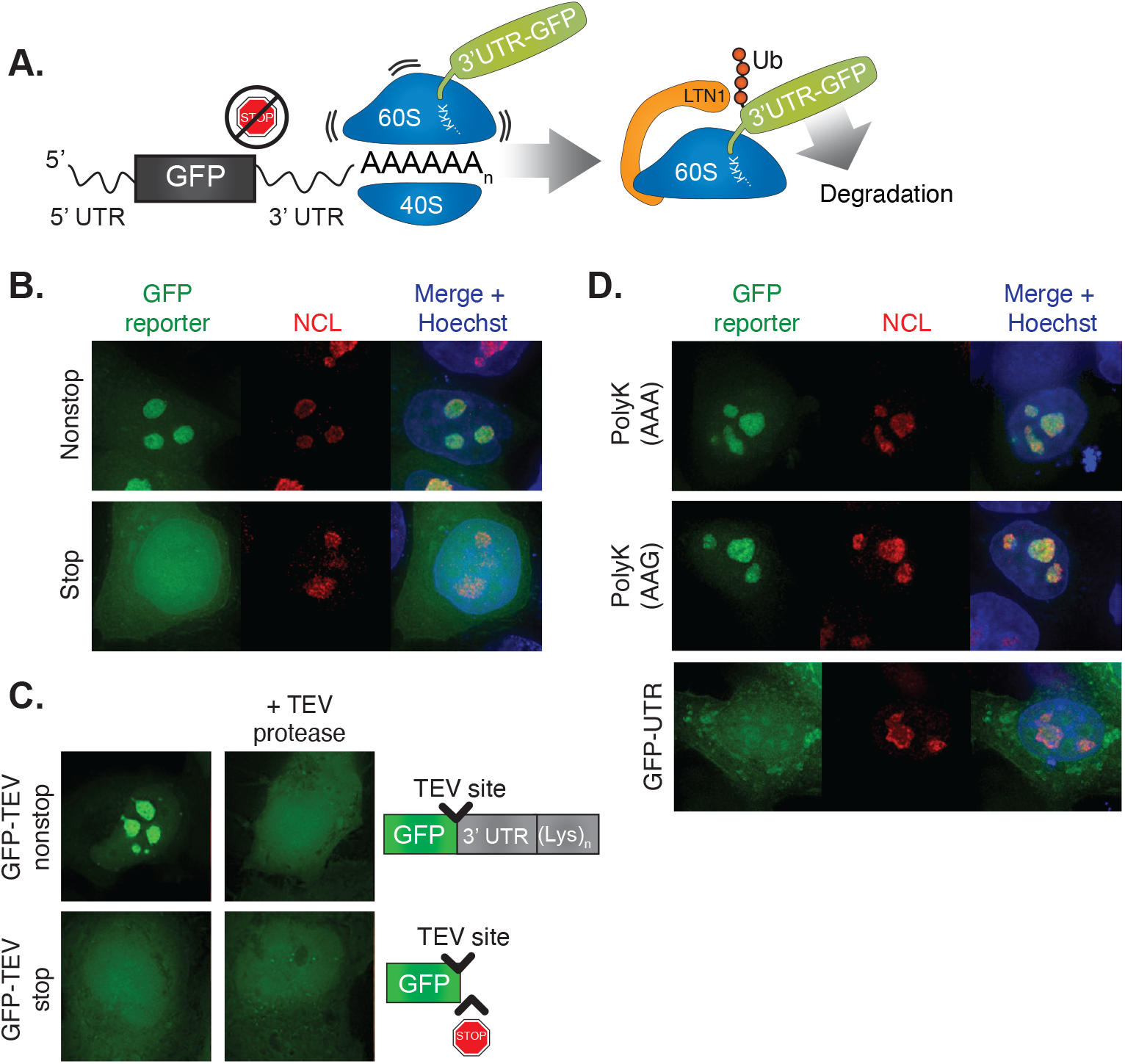
PolyK tracts drive the protein products of non-stop mRNA into nucleoli. (**A**) Schematic of RQC pathway acting on GFP nonstop substrate. Translation proceeds through the UTR into the poly(A) tail, creating a polyK tract. Ribosome stalling then leads to dissociation of the 60S subunit, ubiquitination of the nascent chain by LTN1, and subsequent degradation. (**B**) Localization of GFP-nonstop and GFP-stop. HeLa cells were transfected with GFP reporter constructs, then fixed and immunostained with nucleolin (NCL). (**C**) Effect of C-terminal cleavage on nucleolar localization. HeLa cells were transfected with GFP-TEV-containing constructs (shown in diagram) with or without a plasmid encoding the Tobacco Etch Virus (TEV) protease.

Events that trigger excessive ribosome stalling or compromise RQC have deleterious consequences. On the cellular level, loss of LTN1 function results in proteotoxic stress (Choe et al., 2016; Defenouillère et al., 2016; Yonashiro et al., 2016). On an organismal level, LTN1 hypomorphic mice develop a neurodegenerative disease characterized by progressive impairment of motor and neuronal function (Chu et al., 2009). Similarly, mutation of a tRNA along with loss of GTP Binding Protein 2 (GTPBP2), a protein important for resolving stalled ribosomes, causes neurodegeneration (Ishimura et al., 2014). Yet the mechanisms by which ribosome stalling and defects in co-translational quality control drive neurodegenerative disease remain poorly understood.

To understand the effects of defective proteins produced by stalled translation on cells, we studied the fate of escaped RQC substrates arising from the translation of nonstop mRNA. We found that protein produced from nonstop mRNA accumulated in nucleoli, driven by polybasic tracts from translated poly(A) tails, which served as nucleolar targeting sequences. Nucleolar localization increased the solubility of these defective proteins, potentially providing a protective effect to cells, yet also altered nucleoli. Our study elucidates how failures in co-translational protein quality control can disrupt distal cellular processes and may help define the pathology of neurodegenerative disease.

## Results

### Translation of the poly(A) tail localizes nonstop substrates to nucleoli

To track the fate of proteins produced during stalled translation, we exogenously expressed GFP from a cistron lacking a stop codon (GFP-nonstop) in HeLa cells. This reporter was engineered with the commonly used bGH 3’ UTR because of its well-studied, efficient termination sequence. Mapping the 3’ end of the nonstop transcript confirmed the absence of in-frame stop codons and polyadenylation at the expected site on the reporter mRNA (Fig. S1a). GFP-nonstop protein was stabilized by siRNA knockdown of LTN1, confirming that ribosomes translating GFP-nonstop stalled and triggered RQC (Fig. S1b). Some RQC substrates have been observed to aggregate in yeast (Choe et al., 2016; Defenouillère et al., 2016; Yonashiro et al., 2016). Aggregated proteins can be pathological correlates and possible drivers of disease processes (Aguzzi and O’Connor, 2010; Ross and Poirier, 2004) and thus may underlie stalling and RQC-related neurodegenerative phenotypes. Unexpectedly, we found that GFP-nonstop did not aggregate, but instead localized to discrete structures inside the nucleus (Fig. 1b, top). Staining for nucleolar components revealed that GFP-nonstop localizes to the nucleolus and is distributed throughout all of the nucleolar subcompartments (Fig. 1b, S1c,d), contrasting with the diffuse distribution of GFP-stop (Fig. 1b, bottom).

To verify that the the extra sequence produced from the GFP-nonstop construct (vs. GFP-stop) was required for its nucleolar localization, we removed the C-terminal end of the protein *in vivo* using an engineered TEV cleavage site (GFP-TEV-nonstop) and co-expressed TEV protease. As predicted, this resulted in loss of nucleolar localization (Fig. 1c, top) that was specific to the nonstop construct, as the localization of the diffuse GFP-TEV-stop was unchanged by TEV coexpression (Fig. 1c, bottom).

We sought to determine if an isolated feature of the nonstop protein was sufficient for nucleolar localization. GFP-nonstop translates two additional mRNA regions that differentiate it from GFP-stop: the 3’ UTR and the poly(A) tail. While the UTR can encode many different amino acids, the AAA codons in the poly(A) tail encode for lysine and thus translation of poly(A) tails produces polylysine (polyK) tracts. Previous work has shown that polybasic sequences can act as both nuclear and nucleolar targeting sequences (Scott et al., 2010). Consistent with this, adding a templated, C-terminal polyK tract to GFP was sufficient to cause nucleolar localization (Fig. 1d, top). This nucleolar localization was not codon-specific as polyK tracts encoded by both ‘AAA’ and ‘AAG’ codons yielded comparable localization (Fig. 1d, top and middle). Expression of the UTR sequence without polyK (GFP-UTR) was not sufficient to cause nucleolar-specific localization of the reporter, resulting in localization throughout the cell and the formation of cytoplasmic inclusions. (Fig. 1d, bottom). Thus, we conclude that the polyK tracts of nonstop proteins are necessary and sufficient to localize them to nucleoli.

### Nucleolar targeting increases solubility of amyloidogenic proteins

Nucleolar targeting has been proposed to solubilize misfolded proteins (Frottin et al., 2019; Mediani et al., 2019). To understand the effect of nucleolar localization on our reporters, we took advantage of our observation that GFP-UTR localized to both nucleoli and the cytosol. This allowed us to use an amyloid-specific dye (Amylo-glo) to probe for amyloid state of GFP-UTR in different compartments of the same cell. Strikingly, GFP-UTR stained for Amylo-glo in the cytosol (suggesting the presence of amyloid) but not in the nucleolus (suggesting soluble protein) (Fig. 2a, b). Proteins trapped within aggregates can be less able to exchange with other proteins and thus demonstrate decreased mobility by fluorescent recovery after photobleaching (FRAP) analysis. We found that the perinuclear aggregate fraction of GFP-UTR had dramatically reduced mobility relative to the nucleolar fraction of the protein (Fig. 2c). Collectively, these data suggest that the nucleolar environment solubilized protein that would otherwise have aggregated in the cytosol.

**Figure 2.**
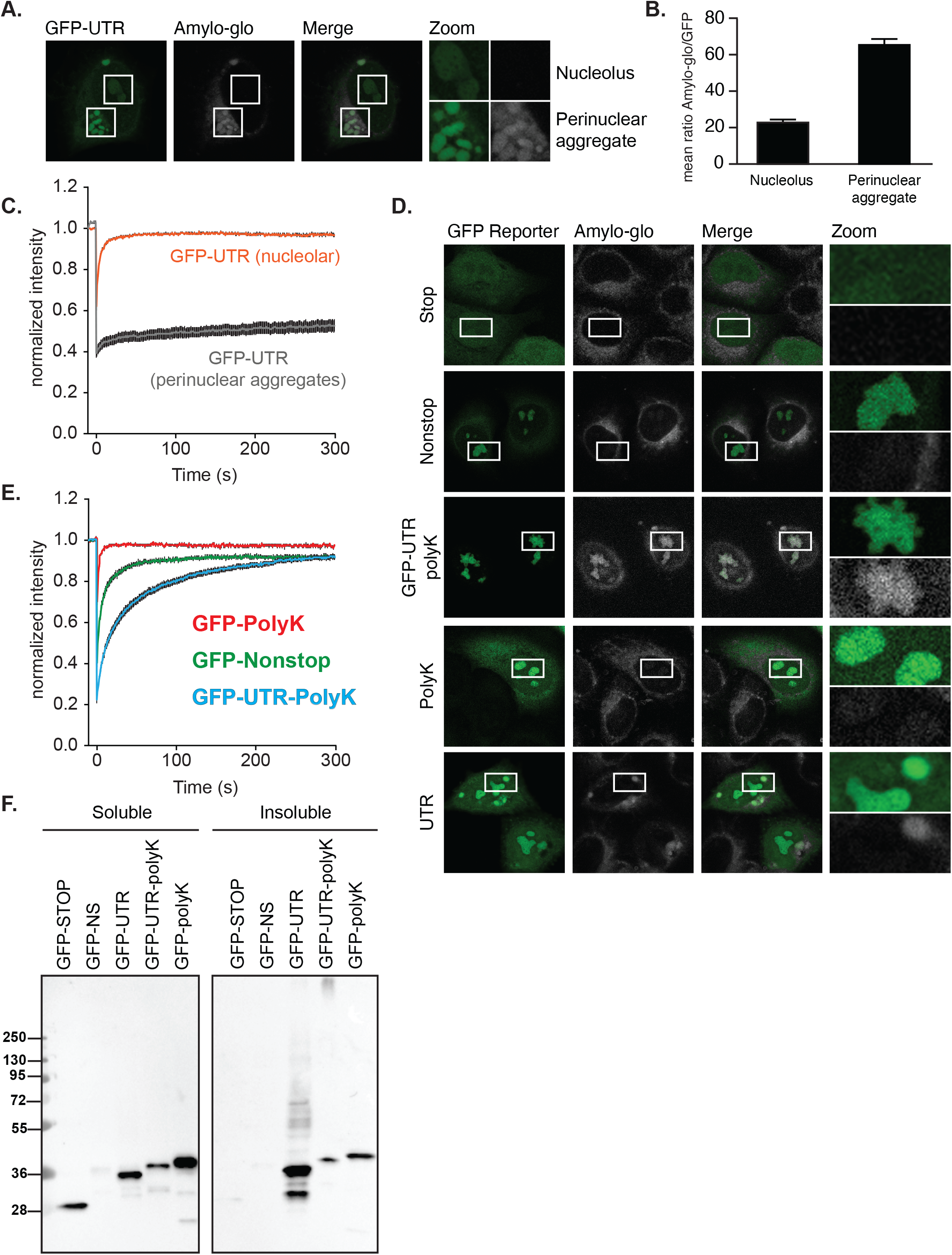
Nucleolar targeting increases solubility of amyloi-dogenic sequences. (**A-B**) 24 hr post-transfection HeLa cells expressing GFP-UTR were assessed for nucleolar and perinuclear amyloid by Amylo-glo staining. Microscopy is quantified in (**B**). The mean Amylo-glo and GFP intensities were measured in 100 ROIs inside nucleoli and 100 perinuclear aggregates, in three independent experiments. *p* = 7.48×10^−27^ (**C**) Fluorescence Recovery After Photobleaching (FRAP) was performed on the nucleolar (orange) and perinuclear (grey) compartments of HeLa cells expressing GFP-UTR. (**D**) Cells expressing GFP constructs for 24 hr were assessed for amyloid by Amylo-glo staining. **(E)** FRAP was performed on the nucleoli of cells expressing the indicated constructs. (**F**) Anti-GFP western blot was performed on solubility-fractionated HeLa cell lysates. Lysate loading volumes were normalized so the soluble and insoluble fractions represent equivalent numbers of cells.

To understand the effect of nucleolar localization of nonstop proteins, HeLa cells were transfected with fluorescent reporters with differing C-termini and stained for amyloid (Fig. 2d). GFP-nonstop-expressing cells stained negatively for nucleolar amyloid, similar to GFP-stop (Fig. 2d, top two panels). However, GFP-nonstop is expressed at relatively low levels due to its mRNA being a constitutive target of nonstop mRNA degradation (Klauer and van Hoof, 2012). To increase the levels of the protein product, we encoded an equivalent sequence to GFP-nonstop on a gene containing a terminal stop codon, GFP-UTR-polyK. GFP-UTR-polyK was expressed at much higher levels and, in contrast to GFP-nonstop, caused strong Amylo-glo staining within nucleoli (Fig. 2d, middle panel). The UTR sequence was essential for this effect, as expression of GFP-polyK resulted in no Amylo-glo staining (Fig. 2d, second to bottom panel), while GFP-UTR produced Amylo-glo positive cytoplasmic aggregates (Fig. 2d, bottom panel).These results suggest that the load of invading proteins determine if the nucleolar environment will be able to solubilize them. Measurements of reporter mobility by FRAP supported this model; GFP-polyK had the highest mobility, GFP-nonstop had a reduced mobile fraction and a somewhat delayed recovery time, and the high-expressing GFP-UTR-polyK had reduced mobile fraction and a further delayed recovery time (Fig. 2e). Using centrifugation to separate soluble and insoluble proteins revealed the same hierarchy of effect; the GFP-stop and most of the GFP-nonstop were found in the soluble fraction (Fig. 2f). By contrast, the majority of the GFP-UTR was insoluble, likely representing the perinuclear aggregates, with the soluble portion being the nucleolar GFP. The GFP-UTR-polyK is proportionately more soluble than GFP-UTR, consistent with an increased solubility in nucleoli versus the cytosol (note that the presence of polylysine also likely contributes to the increased solubility of GFP-UTR-polyK vs. GFP-UTR independent of nucleolar localization). Taken together, these data support a model of enhanced solubility being conferred through localization in the nucleolar compartment and interaction with resident nucleolar proteins.

### LTN1 depletion disrupts nucleoli

We sought to probe the consequences of the influx of endogenous nonstop proteins into nucleoli that is predicted to occur with higher substrate load due to impaired RQC. We blocked RQC by knocking down LTN1, the E3 ligase that marks nonstop proteins for degradation. In cells depleted of LTN1, endogenous nonstop substrates are predicted to escape degradation and build up in the nucleolus. To assay nucleolar integrity, we measured the ability of nucleoli to recover from stress induced by the proteasome inhibitor MG132. MG132 induces nucleolar amyloid body formation (an indicator of nucleolar stress) that can dissolve following the removal of MG132 (Mediani et al., 2019) (Fig. 3a, b). Cells lacking LTN1 had defects dissolving nucleolar amyloid bodies after MG132 removal, suggesting that in the absence of LTN1, nucleoli were more vulnerable to stressors. This is consistent with the model that impaired RQC causes increased levels of nonstop proteins that then invade the nucleolus, interact with resident proteins, and alter nucleolar state.

**Figure 3.**
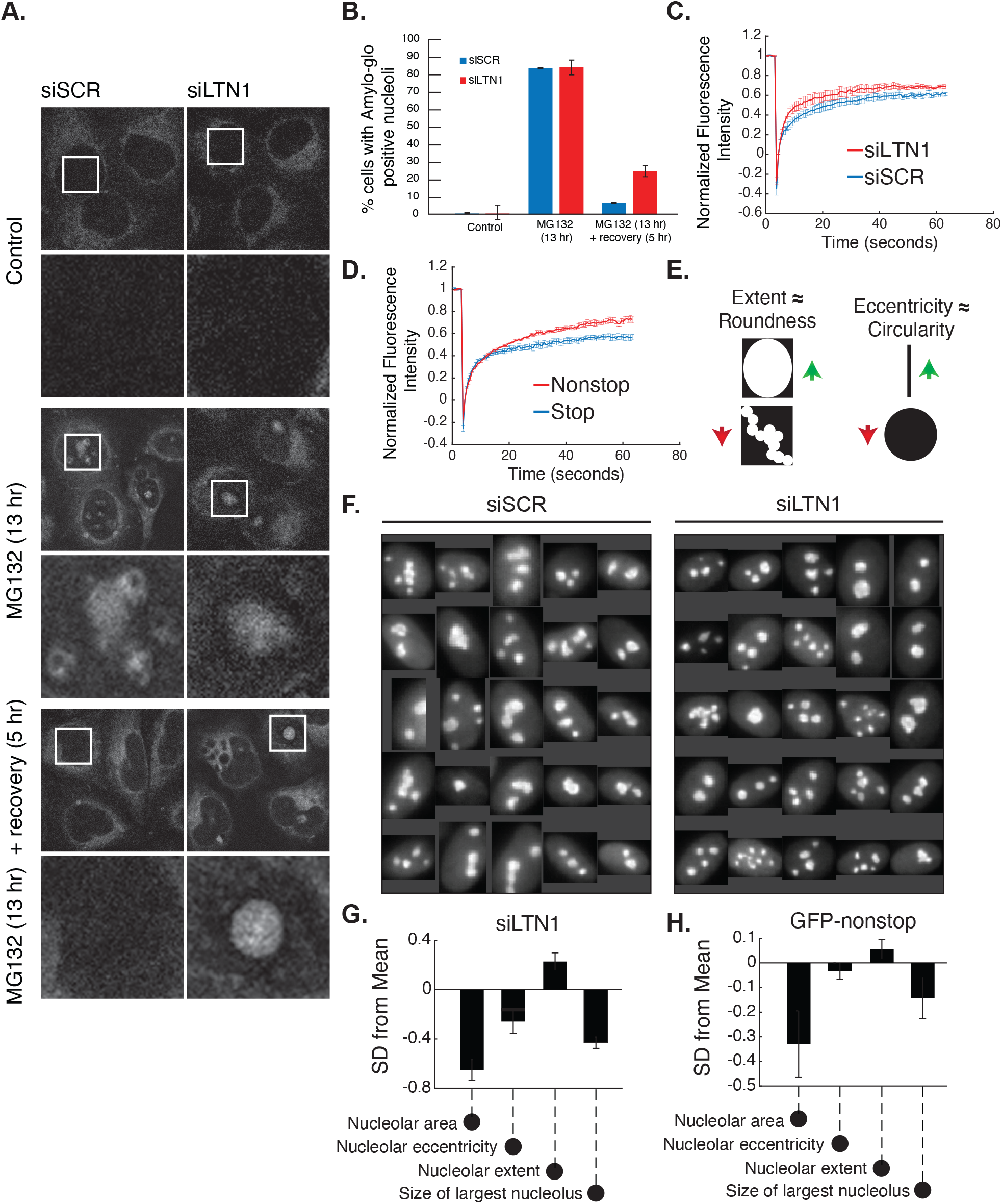
Accumulating nonstop proteins disrupt nucleoli. (**A-B**) siRNA-treated HeLa cells were treated with 10 μM MG132 for 13 hr and allowed to recover for 5 hr. Cells were assessed for amyloid by Amylo-glo staining. Microscopy is quantified in (**B**). (**C-D**) FRAP was performed using GFP-FBL in HeLa cells treated with siLTN1 and siSCR (**C**) or RFP-non-stop and RFP-stop (**D**). Data is presented as means ± the standard error (SE) of three independent experiments (5-20 cells each). (**E-H**) siLTN1-treated and GFP nonstop-expressing HUVEC cells have smaller, rounder nucleoli. Representative images of fixed and NCL-immunostained nucleoli (**F**). Results of automated image analysis quantifying changes in nucleolar morphology of siLTN1 (**G**) and GFP-nonstop (**H**) HUVEC cells. Data are presented as standard deviations (SD) from the mean ± SE of the control distribution [siSCR (**G**) and GFP-stop (**H**)] for >three independent experiments (>500 cells each).

We next explored if interactions with escaped RQC substrates alters nucleolar dynamics. The nucleolus is an archetypal membraneless organelle that is formed at least in part through liquid-liquid phase separation (Feric et al., 2016; Mitrea et al., 2016). Nucleoli are self-organizing and serve to concentrate RNA and proteins into a constrained space, thus enhancing reactions (Lam and Trinkle-Mulcahy, 2015). We hypothesized that an influx of new proteins into nucleoli may thus impact the dynamics of nucleolar-resident proteins. To test this, we examined the mobility of the nucleolar protein fibrillarin (FBL) in cells treated with a non-targeting siRNA control (siSCR) compared with cells treated with siLTN1. Fluorescent recovery after photobleaching (FRAP) of GFP-FBL revealed that the mobile fraction was significantly higher in siLTN1 cells (Fig. 3c), indicating that a larger pool of GFP-FBL is undergoing exchange within nucleoli.

High levels of nonstop protein can exceed the capacity of endogenous LTN1 (Fig. S2a). Thus overexpression of a nonstop reporter serves as an orthogonal method to LTN1 siRNA for increasing the cellular burden of escaped RQC substrates. Exogenous expression of RFP-nonstop but not RFP-stop was sufficient to recapitulate the enhanced mobile fraction of GFP-FBL seen with siLTN1 (Fig. 3d). Taken together, these results suggest that when levels of LTN1 are insufficient to ubiquitylate the pool of polyK-containing nonstop substrates, these substrates enter the nucleolus, interact with resident proteins, and alter nucleolar dynamics.

We next looked to assess changes in nucleolar morphology, as it provides a visible readout of the underlying biophysical state of the nucleolus (Brangwynne et al., 2011). Because HeLa are a transformed cell line with heterogeneous nucleoli, we instead used human umbilical vein endothelial cells (HUVEC), which have more homogeneous nucleolar morphology. HUVEC are primary cells that can be cultured *in vitro* and are amenable to high throughput imaging due to their flat, uniform appearance. Cells were immunostained with the nucleolar marker nucleolin and qualitative differences could be observed between nucleolar morphologies, with siLTN1 nucleoli appearing smaller and rounder than siSCR nucleoli (Fig. 3e-g). To quantify these differences, we systematically analyzed cells (>1500 cells/condition) for nucleolar area, extent (~roundness), eccentricity (~circularity), and the size of the largest nucleolus per cell. Quantitative analysis confirmed that siLTN1 nucleoli were smaller, rounder, more circular, and the largest nucleolus per cell was smaller than siSCR nucleoli. Next, we sought to determine if we could recapitulate the altered nucleolar morphology with GFP-nonstop expression alone, as with the nucleolar mobility phenotype (Fig. 3b). Indeed, expression of the GFP-nonstop substrate had similar effects on nucleolar morphology, with a reduced area, greater extent, less eccentricity, and reduced size of the largest nucleolus per cell relative to a GFP-stop control (Fig. 3h). Both the overexpression of GFP-nonstop (which localizes to nucleoli) and knockdown of LTN1 (which stabilizes endogenous nonstop proteins, which may translate through the poly(A) tail and localize to nucleoli) result in an increase in nucleolar-localized proteins. We next sought to determine if the morphological changes were simply from the excess burden of nucleolar proteins. To address this, we expressed GFP-polyK in HUVEC cells and found that they had no significant changes in nucleolar morphology (Fig. S2b). Collectively, both the disruption of nucleolar dynamics and morphology suggest that nucleoli are affected when a cell’s LTN1 capacity is limited due to siRNA knockdown or from nonstop protein over-expression. Furthermore, the change in nucleolar morphology does not occur simply due to the influx of generic protein into nucleoli, as expression of GFP-polyK did not result in the same effect, and thus appears to be caused by specific features of nonstop substrates.

### Altered nucleolar morphology may be explained by increased intermolecular interaction strength

We sought to understand why an increase in nucleolar proteins might alter morphology through computational modeling. The polyK tracts of nonstop proteins have a strongly positive charge, are predicted to be unstructured, and likely have extensive electrostatic interactions with negatively charged nucleolar material (including nucleic acids). Similarly, FBL contains an unstructured N-terminal RGG domain with a high degree of positive charge, such that we speculated that FBL and polyK tailed proteins could interact with the same binding partners. For our simple model, we assumed that FBL and polyK tracts compete for a single substrate, and that the polyK tail is able to outcompete FBL. This results in decreased eccentricity and size of nucleoli due to stronger intermolecular interactions (Fig. 4a, b). As a quantitative illustration, the relationship between polymer interaction strength, eccentricity, asphericity and droplet volume for a simple two-component (polymer and solvent) phase separated droplet is shown in Fig. 4c (see Supplemental Methods for further details). Thus, the morphological changes to nucleoli are consistent with a model in which the partitioning of nonstop proteins into nucleoli strengthens the driving force for phase separation. Intriguingly, while GFP-polyK partitions into the nucleolus, it does not lead to a change in size and eccentricity (Fig. S2b). This suggests that the observed nucleolar changes with GFP-nonstop are not simply due to the interactions that facilitate partitioning into the nucleolus, but that the translated 3’ UTR, which is predicted to be aggregation-prone and is enriched in hydrophobic residues (Fig. S2c), contributes attractive intermolecular interactions.

**Figure 4.**
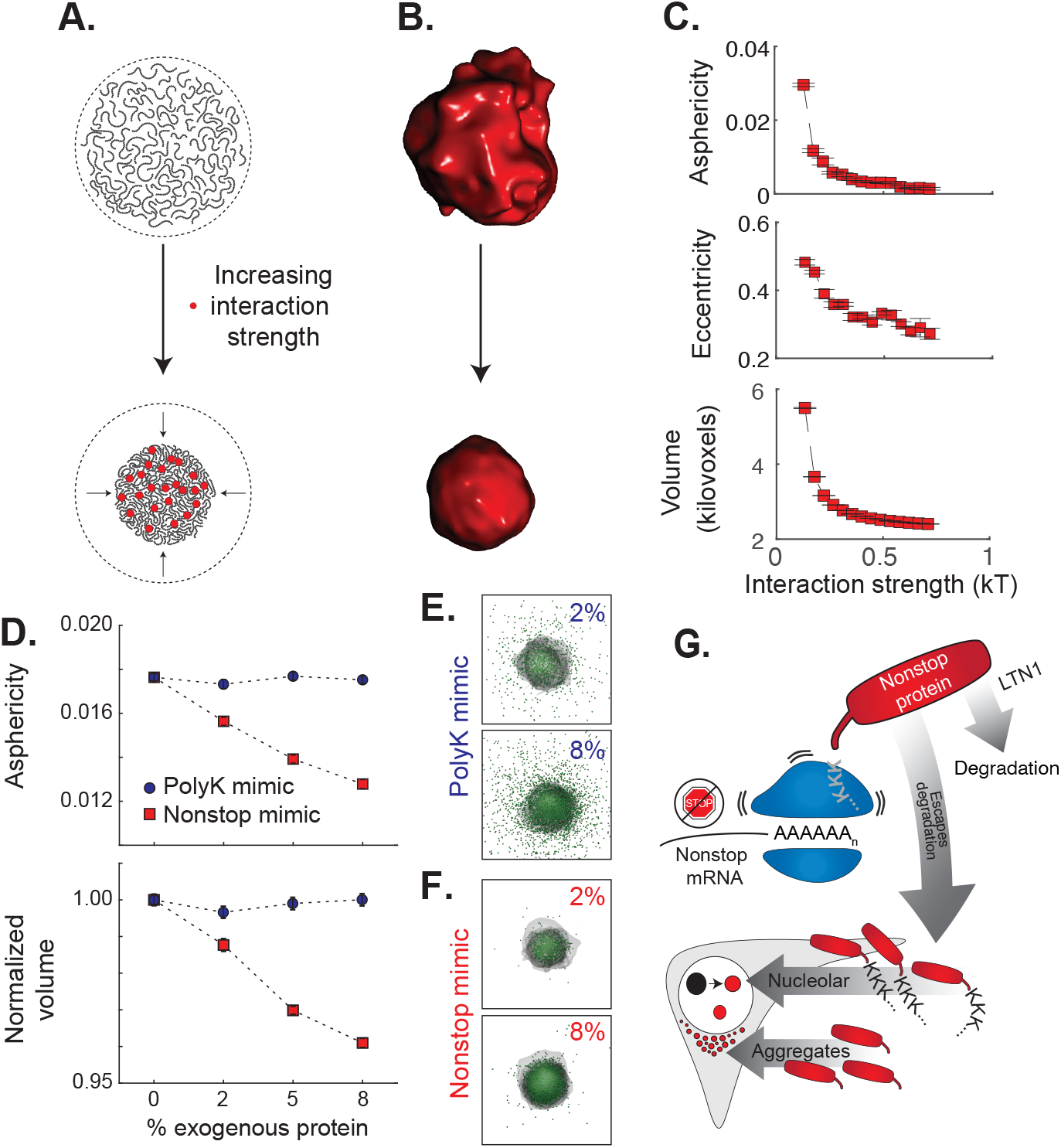
Modeling shows strengthened intermolecular interactions within the nucleolus via nonspecific hydrophobic interactions of nonstop proteins. (**A-B**) Schematic showing how partitioning can drive reduction in droplet volume and increase in asphericity, followed by explicit simulations of a two-component system. (**C**) Assessment of asphericity, eccentricity, and volume for a coarse-grained simulated two-component droplet (polymer and solvent) as a function of the strength of intermolecular interactions. (**D**)Simulation results assessing droplet asphericity and volume as a function of GFP-polyK mimic vs. nonstop mimic. (**E-F**) Snapshots from simulations in panel (**D**). GFP-polyK or nonstop mimic shown as green beads, and the envelope of the nucleolar material is shown as a transparent shell. (**G**) Model for the effect of nonstop proteins on cells: translation of the poly(A) tail of nonstop mRNAs produces a polyK tract. This causes the ribosome to stall, and in the case of insufficient LTN1, cells accumulate nonstop proteins. The polyK tail localizes the nonstop protein to nucleoli, bringing the aggregation-prone, hydrophobic translated 3’ UTR with it. Interactions between nucleolar components and the infiltrating aggregation-prone protein drives increases solubility of defective protein but alteres nucleolar dynamics, morphology, and interferes with the ability to recover from stress.

We next sought to construct a simple biophysical model to explain our observations. We defined a simple single-bead-per-protein lattice-based model in which GFP-polyK or nonstop proteins partition into a phase separated droplet consisting of ‘nucleolar material’. We assume the polyK motif present in both proteins interacts equivalently strongly with nucleolar material and able to outcompete an endogenous nucleolar components (e.g. FBL). A model in which nonstop protein outcompetes FBL is also consistent with the higher FBL mobile fraction observed from FRAP experiments (Fig. 2b). GFP-nonstop may engage in additional non-specific interactions with the nucleolar material driven by hydrophobic residues from the translated 3’ UTR which are absent in GFP-polyK. According to this model, GFP-polyK robustly partitions into the droplet but there are no concentration-dependent changes to droplet morphology (Fig. 4d, e). In contrast, GFP-nonstop protein displays enhanced partitioning and concentration-dependent morphological changes, with droplets becoming smaller and rounder with increasing concentrations of GFP-nonstop (Fig. 4d, f).

While this is a deliberately simple model that does not exclude other possible interpretations, it explains our experimental results. If the translated 3’ UTR engages in strong non-specific interactions with nucleolar components, the gross morphological changes predicted by the model are consistent with our experimentally observed results, where nucleoli become smaller and rounder with LTN1 knockdown or expression of GFP-nonstop.

## Discussion

In this work, we examine how cells are affected when stalled proteins escape co-translational quality control. We find that the protein products of nonstop mRNAs accumulate in nucleoli via polyK tracts produced by translation of poly(A) tails. We posit that the exposed hydrophobic residues of misfolded regions of defective proteins interact with resident nucleolar proteins. This increases the solubility of nonstop proteins but disrupts nucleolar dynamics and morphology (Fig. 4g).

Because 3’ UTRs did not evolve to be translated, they may not have been subjected to negative selection disfavoring aggregation-prone, pathologic sequences. As with our reporter UTR, translation of the 3’ UTR of some disease associated nonstop transcripts have been shown to unmask aggregation-prone, hydrophobic sequences (Bock et al., 2018; Rebelo et al., 2016). Thus, translated 3’ UTRs arising from stop codon readthrough (Dunn et al., 2013) may generally pose a danger to the cell and increasing their solubility by nucleolar localization may be an important quality control strategy. Similarly, misfolded proteins generated by translation of prematurely polyadenylated mRNA may be solubilized and less toxic to cells within nucleoli.

The interaction of escaped quality control substrates with nucleoli is supported by the finding that model substrates are more soluble in nucleoli than in the cytoplasm. This observation dovetails with recent work showing that importins recruited by nuclear localization sequences act as disaggregases in addition to facilitating transport into the nucleus (Guo et al., 2018). Our data suggest that insoluble substrates containing nucleolar localization sequences are kept from aggregating by virtue of being in the nucleolar environment. In some cases, nucleolar sequestration of defective proteins may have a protective effect. For example, a nucleolar localized version of the ALS-associated TDP25 protein decreases cell death by assuming a non-toxic oligomeric state in nucleoli (Kitamura et al., 2016).

Nucleolar sequestration of aggregation-prone proteins occurs upon stress conditions such as heat shock, acidosis, transcriptional stress and proteasome failure. From nucleoli, misfolded proteins are cleared with time by the cell’s protein quality control systems. As such, nucleoli act as reservoirs to prevent irreversible protein aggregation upon stress (Audas et al., 2016; Frottin et al., 2019; Mediani et al., 2019). However, nucleolar sequestration of aggregation-prone proteins is not without consequences; while a causative link has not been conclusively shown, previous work has indicated that disruption of nucleoli may have deleterious consequences (Boulon et al., 2010). Defective clearance of misfolded proteins from nucleoli due to protein quality control impairment leads to irreversible amyloidogenesis and decreases cell viability (Audas et al., 2016; Frottin et al., 2019; Mediani et al., 2019). This has important implications in human disease, as accumulation of aberrant disease-linked proteins such as arginine-rich C9orf72 dipeptide repeats disrupt nucleoli and are causative for Amyotrophic Lateral Sclerosis (ALS) and Frontotemporal Degeneration (FTD) (Lee et al., 2016) and have been associated with neurodegeneration (Frottin et al., 2019; Hernández-Ortega et al., 2016; Mediani et al., 2019; Parlato and Kreiner, 2013). Future work will need to address to what extent nucleolar accumulation of nonstop mRNA protein products contributes to nucleolar dysfunction in ALS/FTD and its role in neurodegeneration caused by a hypomorphic LTN1 allele (Chu et al., 2009).

Remarkably, we found that despite altering nucleolar composition, mobility, and morphology, nucleoli were able to create normal levels of ribosomes and support normal growth rates (Fig. S3a-d). This implies that nucleolar function is largely robust to changes in cellular conditions yet that nucleolar *state* is an early indicator of cellular stress below toxic levels. Nucleolar morphology is currently used as a biomarker for cancer (Derenzini et al., 2009) and we speculate that similar approaches may be useful for detecting stresses relevant to neurodegenerative disease. Our study elucidates how failures in co-translational protein quality control can disrupt distal cellular processes and may help understand the pathological drivers of neurodegenerative disease.

## Materials and methods

### Plasmid construction

The GFP nonstop and stop reporters were constructed by Gibson cloning (Gibson Assembly Master Mix, NEB) eGFP with or without a terminal stop codon into the pCMV AAV backbone which was used as an expression vector and used to make AAV. Stbl3 *E.coli* (Invitrogen) was used for all pCMV AAV vector cloning.The in-frame stop codon in the BGH 3’ UTR was removed to allow for readthrough into the poly(A) tail. The GFP polyK (AAA), polyK (AAG),, and GFP-TEV constructs were constructed by inserting the new sequence with C-terminal stop codons onto the reverse primer, amplifying the GFP nonstop and reinserting into the pCMV AAV backbone with the stop-removed 3’ UTR. Note that if not specified in the text, GFP-polyK is encoded by ‘AAA’ codons. GFP UTR was constructed by amplifying the sequence from GFP-nonstop and using Gateway cloning to insert the sequence with a stop codon before the polyadenylation signal sequence into the pCMV AAV backbone with the no-stop 3’ UTR. The GFP-UTR-polyK construct was made by adding the new sequence to a reverse primer, amplifying from the GFP UTR construct, and inserting into the pCMV AAV backbone with the no-stop 3’ UTR. All constructs were confirmed by sequencing. pEGFP-C1-Fibrillarin was a gift from Sui Huang (Addgene plasmid #26673; http://n2t.net/addgene:26673; RRID: Addgene_26673) (Chen and Huang, 2001).

### Cell culture and transfections

HeLa cells (ATCC) were maintained in Dulbecco’s modified Eagle medium (DMEM; Invitrogen) supplemented with 10% FBS (Sigma) and penicillin-streptomycin (Gemini Bio-Products). The HUVEC human umbilical vein endothelial cell line (ATCC) was grown on collagen-coated vessels and maintained in EGM media supplemented with 2% FBS, hydrocortisone, hFGF-B, VEGF, R3-IGF-1, ascorbic acid, hEGF, and heparin (EGM-2 BulletKit; Lonza). Following the manufacturer’s protocol, Polyjet (SignaGen Laboratories) was used to transfect HeLa cells.

### *In vivo* TEV protease digestion

Constructs containing the TEV cut site were co-transfected with pcDNA3.1 TEV protease into HeLa cells. Cells were fixed and immunostained 24 hr post-transfection. pcDNA3.1 TEV (full-length) was a gift from Xiaokun Shu (Addgene plasmid #64276; http://n2t.net/addgene:64276; RRID: Addgene_64276)(To et al., 2015).

### siRNA knockdowns

Lipofectamine RNAiMAX (Invitrogen) was used for siRNA transfection of HUVEC cells according to the manufacturer’s protocol. siRNA transfections were done every third day with assays performed on day five. Knockdown was confirmed by qPCR and/or by flow cytometry (assaying for siLTN1 stabilization of the RFP-T2A-GFP nonstop reporter).

### Amylo-glo staining

HeLa cells seeded on non-coated glass coverslips were transfected with cDNAs encoding GFP-stop, GFP-nonstop, GFP-UTR, GFP-polyK, and GFP-UTR-polyK using Lipofectamine 2000 (Thermo Scientific) according to the manufacturer’s instructions. 24 hr later, cells were washed with cold PBS, fixed for 9 minutes at room temperature using 3.7% formaldehyde in PBS and permeabilized with 0.2% Triton X-100 inPBS. Cells were then stained for 15 min at room temperature with Amylo-glo (1X) in 0.9% NaCl and washed for 5 min with 0.9% NaCl. Cells were immediately analyzed by confocal microscopy.

For experiments in LTN1-depleted cells, HeLa cells seeded on non-coated glass coverslips were lipofected with either siGENOME Non-Targeting siRNA (siRNA CTL) or with a SMARTpool ON-TARGETplus LTN1 siRNA (L-006968-00-0005, Dharmacon). 48 hr post-transfection, cells were either left untreated or treated with 10 µM MG132 for 13 hr, followed by recovery in drug-free medium for 5 hr. Cells were then processed for Amylo-glo staining as described above.

### Immunostaining

On the fifth day following the start of knockdown, the day after reporter transfection, or the second day after AAV reporter infection, HUVEC cells were washed once with PBS (Invitrogen) and fixed for 15 minutes with 4% formaldehyde (Ted Pella). After two PBS washes, cells were permeabilized and blocked with 10% FBS, 1% BSA, and 0.1% Triton X-100 (Sigma-Aldrich) for 30 min. Then, mouse α-nucleolin (C23) antibody (clone MS-3; Santa Cruz Biotechnologies) diluted in permeabilization/block buffer (1:5000) was added for 2-4 hr at room temperature. Following three 5 min PBS washes, goat α-mouse IgG secondary antibody (DyLight 650; Thermo Fisher Scientific) diluted in permeabilization/block buffer (1:1000) was incubated for 1-2 hr at room temperature. After three PBS washes, nuclei were stained with Hoescht. Other antibodies used for immunostaining and any procedural modifications are listed below: mouse α-UBF (1:100, Santa Cruz Biotechnologies) - cells were fixed with ice-cold methanol for 5 min, then blocked and stained with primary and secondary antibody in 10% FBS and 1% BSA in PBS; rabbit α-FBL (1:500, One World Labs); and rabbit α-NPM1 (1:1000, Santa Cruz Biotechnologies).

### Fractionation of NP-40 soluble and insoluble proteins

24 hr after transfection, cells were harvested in a buffer containing 1% NP-40 (50 mM Tris-HCl, pH 7.4, 150 mM NaCl, 0.25% deoxycholic acid, 1% NP-40, and 1 mM EDTA) and passed through a 26G needle 3 times. Cells were lysed on ice for 10 min, then spun at 10,000 x g at 4°C for 10 min. The supernatant was collected as NP-40 soluble fraction, while the pellet (NP-40 insoluble fraction) was resuspended with 2% SDS Laemmli buffer. Protein fractions were boiled for 3 minutes at 100°C, reduced with β-mercaptoethanol and separated by SDS-PAGE, followed by western blot using an anti-GFP antibody (632381, Clontech).

### High-throughput nucleolar morphology microscopy

HUVEC cells were infected with AAV (MOI of 10) expressing reporter constructs for two days or siRNA knockdown was performed for 5 days. Cells were then fixed and immunostained as described in the Immunostaining section. Images were acquired with MicroManager v1.4 software package(Edelstein et al., 2010, 2014) on a Nikon Eclipse TiE inverted fluorescence microscope with a 40x (NA 0.95) air objective (Nikon Instruments) equipped with a CCD camera (Andor Technologies). Data were collected from a minimum of three independent experiments. For each experiment, at least 500 cells were analyzed for each condition. The Matlab software platform (MathWorks) was used for high-throughput image analysis. First, a mask covering the nucleus was constructed from the Hoescht image. Very large (cells touching) or very small (debris or dividing cells) nuclei were not included in the analysis. Next, the program looked within the nuclear area defined by the Hoescht mask in the nucleolin channel to identify nucleoli. From here, parameters such as area, perimeter, extent (roundness), and eccentricity (circularity) were calculated for the different conditions.

### Other fixed cell microscopy

Images were acquired on a CoolSnap HQ CCD camera (Photometrics) with a DeltaVision Core deconvolution microscope using 0.2 µm steps. Pictures shown in Figures 2A, D and 3A were acquired using a Leica SP8 confocal microscope equipped with a 405nm and white light lasers using a 63X oil immersion objective. The Amylo-glo and GFP intensities were calculated in specific ROI using ImageJ (https://imagej.nih.gov/ij) and the mean ratio intensity of Amylo-glo was divided by the mean ratio intensity of GFP. 100 cells in three independent experiments were analyzed.

### Fluorescence recovery after photobleaching (FRAP)

Cells were plated on glass-bottom 96-well plates (Cell-Vis). After 18-24 hr, GFP FBL was transfected, and cells were imaged 24 hr later. FRAP was performed with an inverted Zeiss LSM 880 laser scanning confocal microscope using a 63x oil objective. A small region of interest was bleached by scanning with the 405 nm laser at 100% power. A normalizing, unbleached area as well as a non-fluorescent background region were also collected. Images were captured every 0.663 sec for a total of 100 images. Data was exported from FRAP movies using Fiji (Schindelin et al., 2012) with the Bio-Formats plugin(Linkert et al., 2010). Data was analyzed using the easyFRAP-web single-page application(Koulouras et al., 2018). Full scale normalized data was plotted against time and fit with a double exponential. FRAP measurements on HeLa cells transfected with cDNAs encoding GFP-nonstop, GFP-UTR, GFP-polyK and GFP-UTR-PolyK were performed on a Leica SP8 confocal microscope equipped with a 405nm and white light lasers using a 63X oil immersion objective. A nucleolus was bleached for 1 sec using a laser intensity of 100% at 405nm. Recovery was recorded for 120 time points after bleaching (120 sec). Analysis of the recovery curves was carried out using a custom written FIJI/ImageJ routine. The equation used for FRAP analysis is as follows ((Ibleach - Ibackground)/(Ibleach(t0) - Ibackground(to)))/((Itotal - Ibackground)/(Itotal(t0) - Ibackground(to))), where Itotal is the fluorescence intensity of the entire cellular structure, Ibleach represents the fluorescence intensity in the bleach area, and Ibackground the background of the camera offset. Fluorescent density analysis was performed using FIJI/ImageJ and selecting specific region of interest (ROI). When necessary, image drift correction was applied using StackReg plug‐in function of the FIJI software suite prior to FRAP analysis. FRAP curves were averaged to obtain the mean and standard deviation.

### Flow cytometry

HeLa cells were trypsinized and resuspended in serum-free media supplemented with 5 mM EDTA prior to measurement on an Accuri C6 plus cytometer equipped with 488 and 640 nm lasers (BD Biosciences). Untransfected cells were gated out during analysis.

### AAV production and infections

293T cells were transfected with adenovirus helper, AAV helper and pAAV CMV GFP constructs using HBS-calcium chloride transfection. Adherent and floating cells were collected after 3 days and freeze-thawed three times, treated with Benzonase for 30 min at 37°C, and supernatants were collected after spinning for 13,500 rpm for 30 min at 4°C.

### Hydropathicity assessment

The translated sequence of the GFP-nonstop 3’UTR was run through the ExPASy ProtScale amino acid calculator for hydrophobicity (Gasteiger et al., 2005) using the Kyte and Doolittle scale (Kyte and Doolittle, 1982). The average hydrophobicity value of the translated human proteome was calculated based on calculated amino acid abundances (Nakamura et al., 2000).

### Simulations

Simulations were performed using a coarse-grained model in which proteins and nucleolar material are represented as one or more beads on a cubic lattice. A nearest-neighbour lattice interaction potential was used, and simulations were run under conditions in which a single phase-separated droplet is formed and stable at equilibrium. Further details can be found in the Supplemental Methods.

## Supplemental methods

### Plasmid construction

The RFP-T2A-GFP nonstop and stop reporters were constructed by Gibson cloning the RFP (mKate2), T2A, GFP (+/− stop codon), and pCDNA4.TO together. This resulted in a single mRNA capable of producing two physically separate proteins (RFP and GFP) (Kim et al., 2011). The RFP-T2A-GFP GFP polyK(AAA) and (AAG) reporters were generated by restriction enzyme cloning. The region between the 5’ SnaBI (Eco1051) and 3’ EcoRI sites in the reporters were PCR amplified and the additional sequence (with a stop codon) was added on the reverse primer. This fragment was then ligated into the RFP-T2A-GFP pCDNA4.TO backbone. All constructs were confirmed by sequencing.

### Flow cytometry

For the dual color RFP-T2A-GFP reporter, differences in plasmid expression levels were accounted for by normalizing nonstop GFP levels to the RFP control.

### EU staining

HUVEC cells were incubated with EGM-2 media containing 100 μM 5-ethynyl uridine (EU) for 25 min. Cells were then fixed with 4% paraformaldehyde for 15 min and permeabilized for 30 min with 0.1% Triton X-100 (Sigma-Aldrich) in PBS. Alexa-Fluor 594 azide was used to label nascent RNAs following the Click-iT RNA Imaging Kit (C10330, Invitrogen) protocol.

### Luciferase viability assay

HeLa cells were knocked down with siRNA for 5 days as described. On day 5, cells were processed following the ONE-Glo + Tox cell viability assay (Promega).

### Bioanalyzer

Total RNA was isolated from siRNA-treated cells using RNA-Bee (Tel-Test). RNA was analyzed using a 2100 Bioanalyzer (Agilent) with the eukaryotic total RNA pico chip (Agilent).

### Simulations

Simulations were performed on a cubic 3D lattice. Proteins are represented either as single beads or as polymers of beads. Each bead only interacts with neighboring beads via a simple intermolecular potential. The position of beads and polymers on the lattice are evolved using a series of Monte Carlo moves evaluated by the Metropolis criterion that include molecular translation/rotation, chain pivoting, and cluster translation/rotation. All simulations were run for ~3×10^8^ moves. In all simulations analyzed, a single droplet is formed and remains stable and at-equilibrium for the duration of the simulation. Volume is calculated through the projection of a spherical envelope over the droplet. Eccentricity provides a measure of deviation from circularity and is scaled between 0 (circle) and 1 (extended oblong) and is calculated by taking 2D snapshots and performing image analysis of the resulting images using MATLAB (MathWorks). Asphericity provides a measure of deviation from a perfect sphere and is scaled between 0 (sphere) and 1 (prolate ellipsoid) and is calculated directly from the droplet gyration tensor(Holehouse et al., 2015). Error is calculated as the standard error of the mean. For two-component simulations (Fig. 4b, c) 500 5-bead homopolymers were used. The interaction strength between beads were titrated under a regime in which the polymers undergo phase separation. Analysis was performed over the second half of the simulation. Five independent simulations were performed per interaction strength.

For multi-component simulations (Fig. 4d, e) proteins (GFP-polyK and nonstop protein) are represented as single beads. Nucleolar material is represented by two different species; a 3-bead homopolymer (nucleolar RNA) and a single bead (FBL). All simulations use 300 nucleolar RNA polymers (900 beads) and a total number of 600 additional proteins. In the absence of GFP-polyK or nonstop protein, nucleolar RNA and FBL undergo phase separation to form a stable two-phase system. To recapitulate the ability of GFP-polyK and nonstop protein to outcompete FBL, we assume these components replace an equivalent number of FBL in our system. This provides a simple manifestation of mutually exclusive interactions, although this mode can also be achieved directly using a more complex interaction scheme that yields equivalent results. As the number of GFP-polyK or nonstop protein increase, these components partition into the nucleolus. However, only the nonstop protein engages in additional interactions, leading to the decrease in droplet volume and increase in asphericity. For all conditions, fifty independent simulations were performed.

While this is the simplest schema, we could construct to provide a physical model of our system, we do not mean to suggest these simulations reflect the full complexity of the underlying observed phenomena within the nucleolus. As an example, given our experiments demonstrate that the partitioning of nonstop protein in the nucleolus reduces nucleolar volume, a possible anticipated corollary is that there is a corresponding increase in nucleolar density. From fluorescence studies we see no indication of such an increase in density (minimal change in brightness), a result consistent with a model in which nonstop protein outcompetes one or more components in the nucleolus. However, we cannot exclude the possibility that changes within the nucleolar environment alter the manner in which fluorescence should be interpreted (a topic of ongoing study by a number of groups). In short, given the complexity of the system, there are a number of unknown variables we have chosen not to consider when drawing mechanistic conclusions, despite the fact that the simplest explanation are broadly consistent with our model. Instead, we present our simulation results as a simple, physical model of the emergent behaviour expected upon the partitioning of an attractive species into a phase separated body that outcompetes an endogenous component, a model that yields results consistent with both the nucleolar dysregulation and the cytoplasmic aggregation of the 3’ UTR.

## Acknowledgements

We thank the members of the Brandman laboratory for helpful discussions and commentary on the manuscript. Additionally, we acknowledge the members of the Julie Theriot Lab (Stanford/University of Washington) and Aaron Straight (Stanford) labs for use of microscopes, reagents, and expertise. We also thank people from the CIGS microscopy facility (University of Modena and Reggio Emilia) for technical support. This research was supported by a Stanford Dean’s Postdoctoral Fellowship to ZHD as well as a grant (1R01GM115968) from the National Institutes of Health to OB. This work was performed while ASH was a postdoctoral fellow in the laboratory of Dr. Rohit V. Pappu (RVP) at Washington University in St. Louis and supported by the Human Frontiers Science Program (grant RGP0034/2017 to RVP). S.C. acknowledges funding from AriSLA Foundation (MLOpathy); Cariplo Foundation (Rif. 2014-0703); MAECI (Dissolve_ALS); MIUR (Departments of excellence 2018-2022; E91I18001480001). S.C. and S.A. are grateful to EU Joint Programme - Neurodegenerative Disease Research (JPND) project. The project is supported through funding organisations under the aegis of JPND (http://www.neurodegenerationresearch.eu/). This project has received funding from the European Union’s Horizon 2020 Research and Innovation Programme under grant agreement No 643417.

---

**Figure S1.**
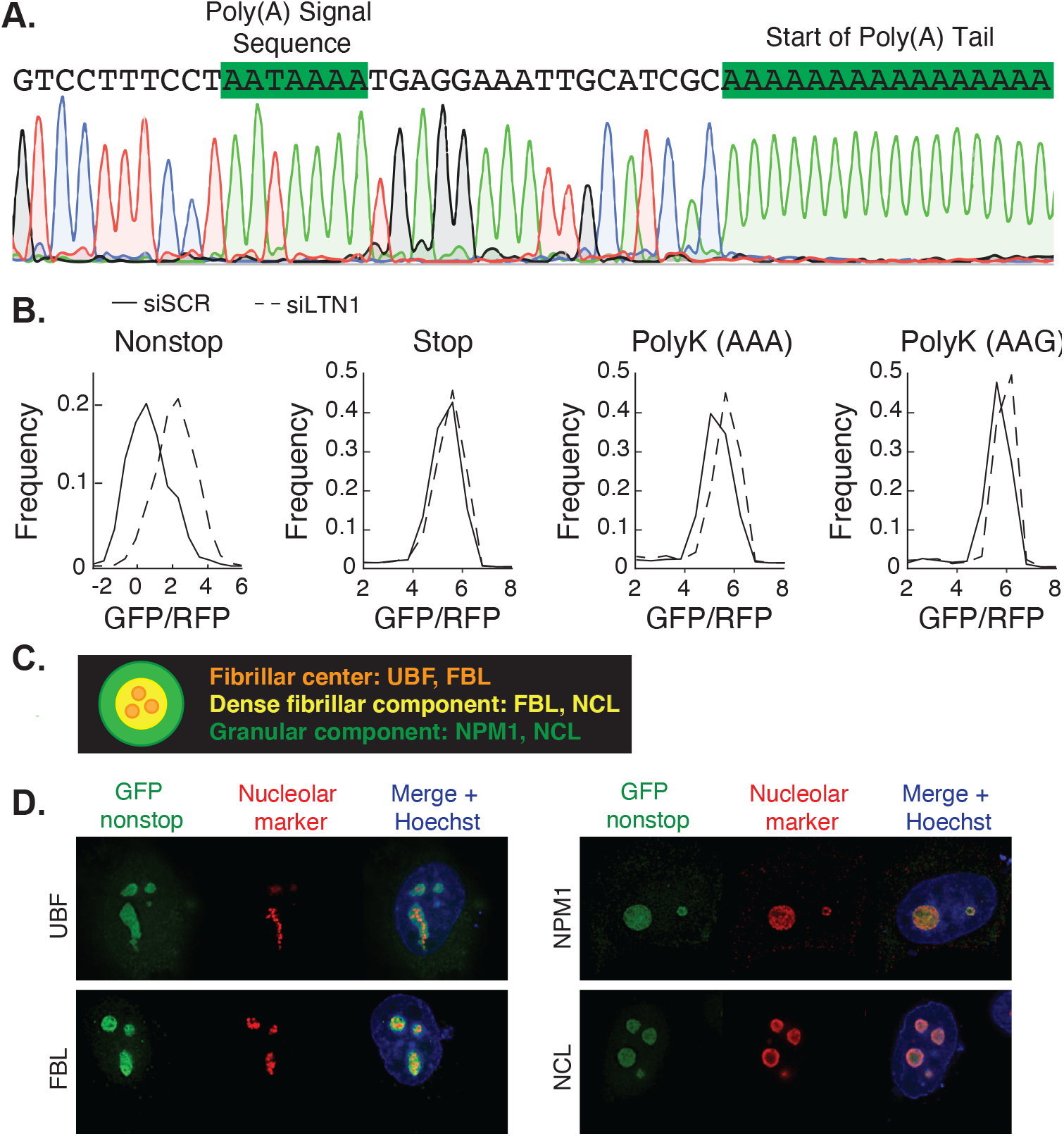
GFP-nonstop has no in-frame stop codons, is degraded by LTN1, and localizes to nucleoli. (**A**) A representative sequencing chromatogram of 3’ Rapid Amplification of cDNA Ends (RACE) analysis showing the site of transcript polyadenylation. (**B**) As an expression control, RFP was encoded upstream of the GFP sequence in the same open reading frame and separated by a P2A peptide bond skipping sequence (Kim et al., 2011). The translated RFP is released before the ribosome encounters the 3’ portion of the mRNA containing the GFP-nonstop sequence, the 3’ untranslated region (UTR) and poly(A) tail. This allows changes in GFP to be internally normalized to RFP, facilitating comparisons in GFP degradation between different conditions and reporters. (**C-D**) Measurements of GFP-nonstop localization within nucleolar subcomponents as assayed by co-localization with fibrillar center, dense fibrillar, and granular component markers.

**Figure S2.**
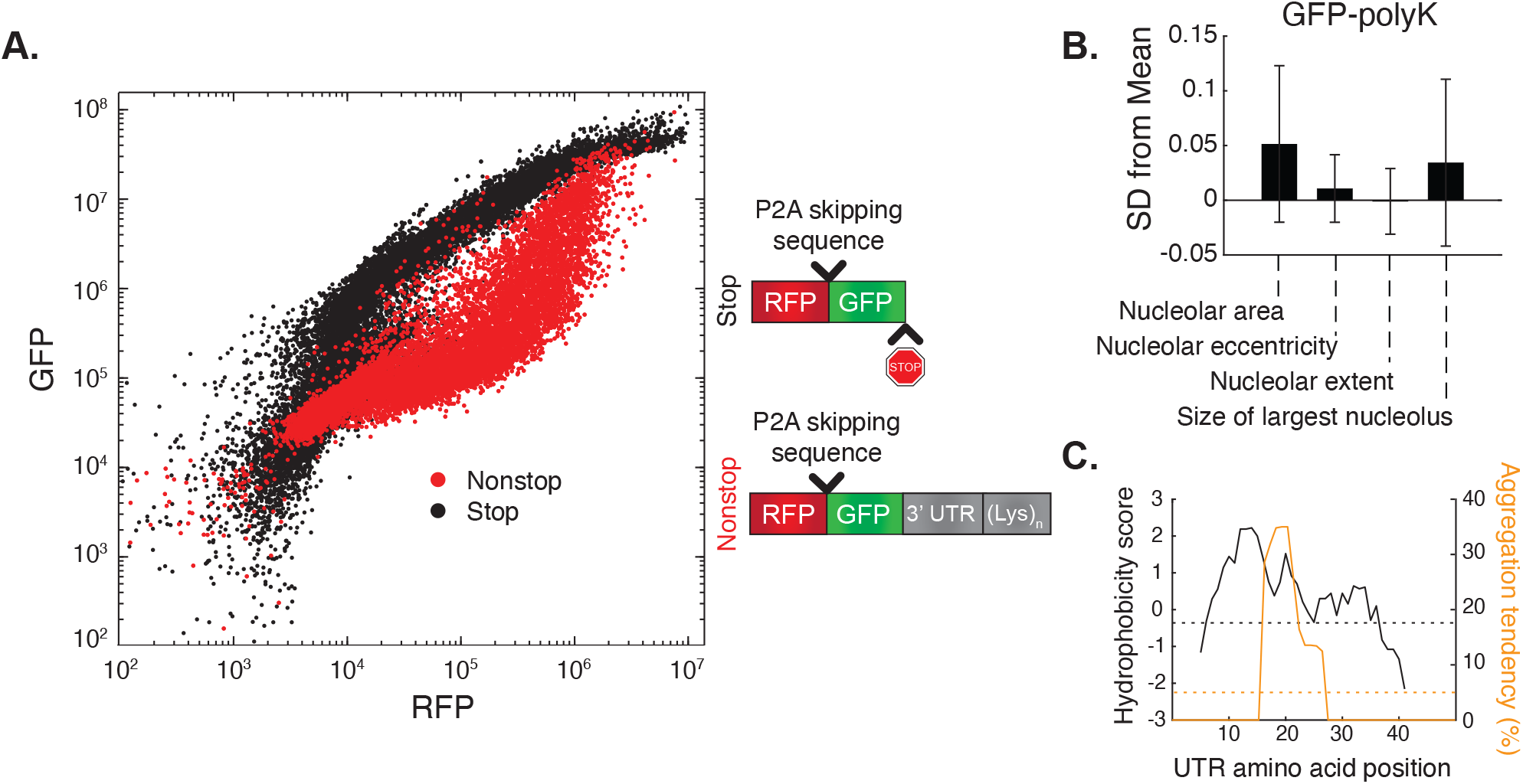
GFP-polyK does not alter nucleolar morphology and high levels of GFP-nonstop can saturate endogenous LTN1. (**A**) Flow cytometry data for HeLa cells transfected with the indicated two-color reporter. (**B**) Quantified changes in nucleolar morphology of GFP-polyK (normalized to GFP-stop). Data are presented as standard deviations (SD) from the mean ± SE of the control distribution (GFP-stop) for >three independent experiments (>500 cells each). (**C**) Hydrophobicity score (black) of the translated 3’ UTR protein sequence, represented as 9 residue moving averages, where a higher score indicates greater hydrophobicity. The dashed black line represents the average hydrophobicity score of the human proteome based on the overall abundance of each amino acid (−0.36). Predicted aggregation tendency of the 3’ UTR is depicted in orange. The dashed orange line indicates the threshold for a potentially aggregating segment (5%).

**Figure S3.**
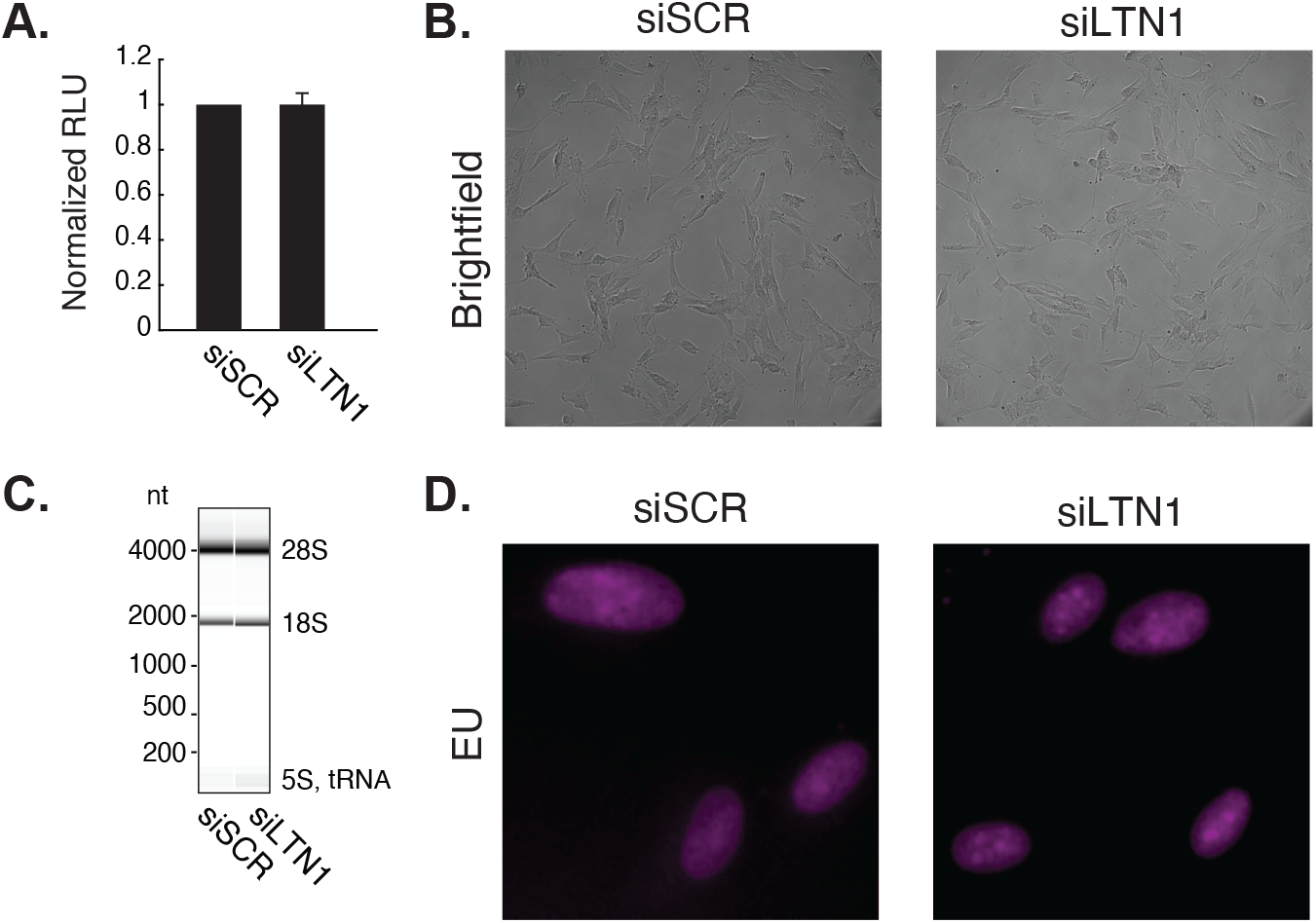
LTN1 knockdown does not affect viability or growth. (**A**) Luciferase cell viability assay of siRNA-treated cells. Data are presented as means ± SE. (**B**) Representative brightfield images of live (unfixed) siRNA-treated cells. (**C**) Representative Bioanalyzer traces of total RNA isolated from siRNA-treated cells. (**D**) 5-ethynyl uridine (EU) incorporation and staining indicating newly synthesized RNA.

